# Recognition of *Chlamydia trachomatis* by Toll-Like Receptor 9 is altered during persistence

**DOI:** 10.1101/2024.02.06.579186

**Authors:** Aissata Diallo, Grace Overman, Prakash Sah, George W. Liechti

## Abstract

Toll-like receptor 9 (TLR9) is an innate immune receptor that localizes to endosomes in antigen presenting cells and recognizes single stranded unmethylated CpG sites on bacterial genomic DNA. Previous bioinformatic studies have indicated that the genome of the human pathogen *Chlamydia trachomatis* contains TLR9 stimulatory motifs, and correlative studies have implied a link between human TLR9 (hTLR9) genotype variants and susceptibility to infection. Here we present our evaluation of the stimulatory potential of *C. trachomatis* gDNA and its recognition by hTLR9– and murine TLR9 (mTLR9)-expressing cells. We confirm that hTLR9 colocalizes with chlamydial inclusions in the pro-monocytic cell line, U937. Utilizing HEK293 reporter cell lines, we demonstrate that purified genomic DNA from *C. trachomatis* can stimulate hTLR9 signaling, albeit at lower levels than gDNA prepared from other Gram-negative bacteria. Interestingly, we found that while *C. trachomatis* is capable of signaling through hTLR9 and mTLR9 during live infections in non-phagocytic HEK293 reporter cell lines, signaling only occurs at later developmental time points. Chlamydia-specific induction of hTLR9 is blocked when protein synthesis is inhibited prior to the RB-to-EB conversion and exacerbated by the inhibition of lipooligosaccharide biosynthesis. The induction of aberrance / persistence also significantly alters Chlamydia-specific TLR9 signaling. Our observations support the hypothesis that chlamydial gDNA is released at appreciable levels by the bacterium during the conversion between its replicative and infectious forms and during treatment with antibiotics targeting peptidoglycan assembly.

## INTRODUCTION

*Chlamydia trachomatis* is an obligate intracellular bacterium that utilizes a unique biphasic developmental cycle^1^. The pathogen has three morphological forms; an Elementary Body (EB) that is infectious, but non-replicating, a Reticulate Body (RB) that is the replicative but non-infectious, and an aberrant body (AB) that is thought to be a defensive response by the bacterium to extracellular stressors. The microbe spends the majority of its existence in an intracellular pathogenic vesicle called an inclusion, where it is protected from a number of immune responses to bacterial infections^2^.

Chlamydia species are recognized by the innate immune system via Toll-like receptors (TLRs) and NOD-like Receptors (NLRs), which respond to various components of bacterial and viral pathogens^3^. This early interaction initiates a cytokine signaling cascade that results in the recruitment of immune cells to the area in order to fight off the infection and direct a subsequent, cell-mediated adaptive response^4^. TLRs play an important role in numerous infections of the female lower genital tract^5^, and a number of studies have investigated the importance of various immunostimulatory chlamydial components (lipids / lipoproteins / MOMP / HSP60, Lipooligosaccharide; LOS, peptidoglycan) and their cognizant innate immune receptors (TLR2, TLR4, NOD1/2), respectively^3^. The predominantly held view is that TLR2 plays a major role during chlamydial infections, while TLR4 and NOD1/2 are accessory and play a more supportive role^6^. This assessment is based largely on the fact that Chlamydia species produce less immunostimulatory LOS^7–9^ and peptidoglycan^10–14^ than similarly sized Gram-negative bacteria, supporting the hypothesis that these differences represent pathoadaptations by the organism that enable it to subvert the host’s innate immune system^2,15^.

Considerably less is known about the role of the innate immune receptor TLR9 during Chlamydia infections. TLR9 recognizes single stranded, unmethylated cytosine-phosphate-guanosine (CpG) sites on bacterial genomic DNA^16^. Unlike TLR2/4 and NOD1/2, which are broadly expressed by a large number of cell types, TLR9 is expressed almost exclusively in monocytes and dendritic cells^16–18^. In addition to being expressed in only a small subset of cell populations, TLR9 is exclusively found within vesicles associated with the endosomal maturation pathway. When phagocytic cells ingest foreign, extracellular bacteria, TLR9 is translocated to the endosomal compartment^19^ and signals after trafficking to the lysosome where it interacts with the DNA released from bacteria undergoing degradation^20^. While *C. trachomatis* avoids fusing with vesicles in the endosomal maturation pathway in most cell types^21–27^, this is not the case in many immune cells that actively express TLR9^28–31^.

Because many TLR9 stimulatory (and inhibitory) signaling motifs have been identified, it is possible to estimate the stimulatory potential of genomic DNA from any microbe whose genome has been sequenced^32^. *C. trachomatis* stimulatory DNA CpG motifs have been calculated in the low-to-mid range^33,34^, indicating that its gDNA is likely recognizable by TLR9. Correlative studies have found that TLR9 gene polymorphisms are associated with increased risk of cervicitis^35^ and pathology associated with Chlamydia infections in human patients^36^, as well as susceptibility to Chlamydia infection in ruminants^37^. Despite these observations, studies investigating TLR9’s role in responding to Chlamydial infections in animal models have demonstrated little to no significant association with infectivity or disease presentation^34,38,39^, leading us to question the degree to which TLR9 signaling occurs in Chlamydia-infected cells.

Here we present an analysis of the stimulatory potential of gDNA from *C. trachomatis* and through a series of experiments demonstrate its ability to interact with TLR9.

## RESULTS

### Genomic DNA from *C. trachomatis* induces hTLR9 signaling in vitro

We began our study by first confirming that gDNA isolated from *C. trachomatis* was capable of initiating a signaling cascade through TLR9. Utilizing a human TLR9-expression reporter cell line (hTLR9-HEK-Blue) we examined secreted Embryonic Alkaline Phosphatase (SEAP) activity in the supernatants of reporter cells exposed to various concentrations of sonicated gDNA obtained from three different bacterial species; *Escherichia coli* (strain MG1655), *Chlamydia trachomatis* (serovar L2, strain Bu / 434), and *Chlamydia muridarum* (strain Nigg). *E. coli* gDNA was highly stimulatory, *C. muridarum* gDNA exhibited no stimulatory activity, and *C. trachomatis* gDNA was somewhat stimulatory, ∼50x less stimulatory than *E. coli* gDNA **(Fig. 1)**. We took these results as confirmation that gDNA (purified from *C. trachomatis* EBs) can act as a positive stimulatory ligand for hTLR9, but that Chlamydia-specific TLR9 signaling is (as predicted by in silico analysis^33^) not as robust as that of other Gram-negative bacteria.

**Figure 1:**
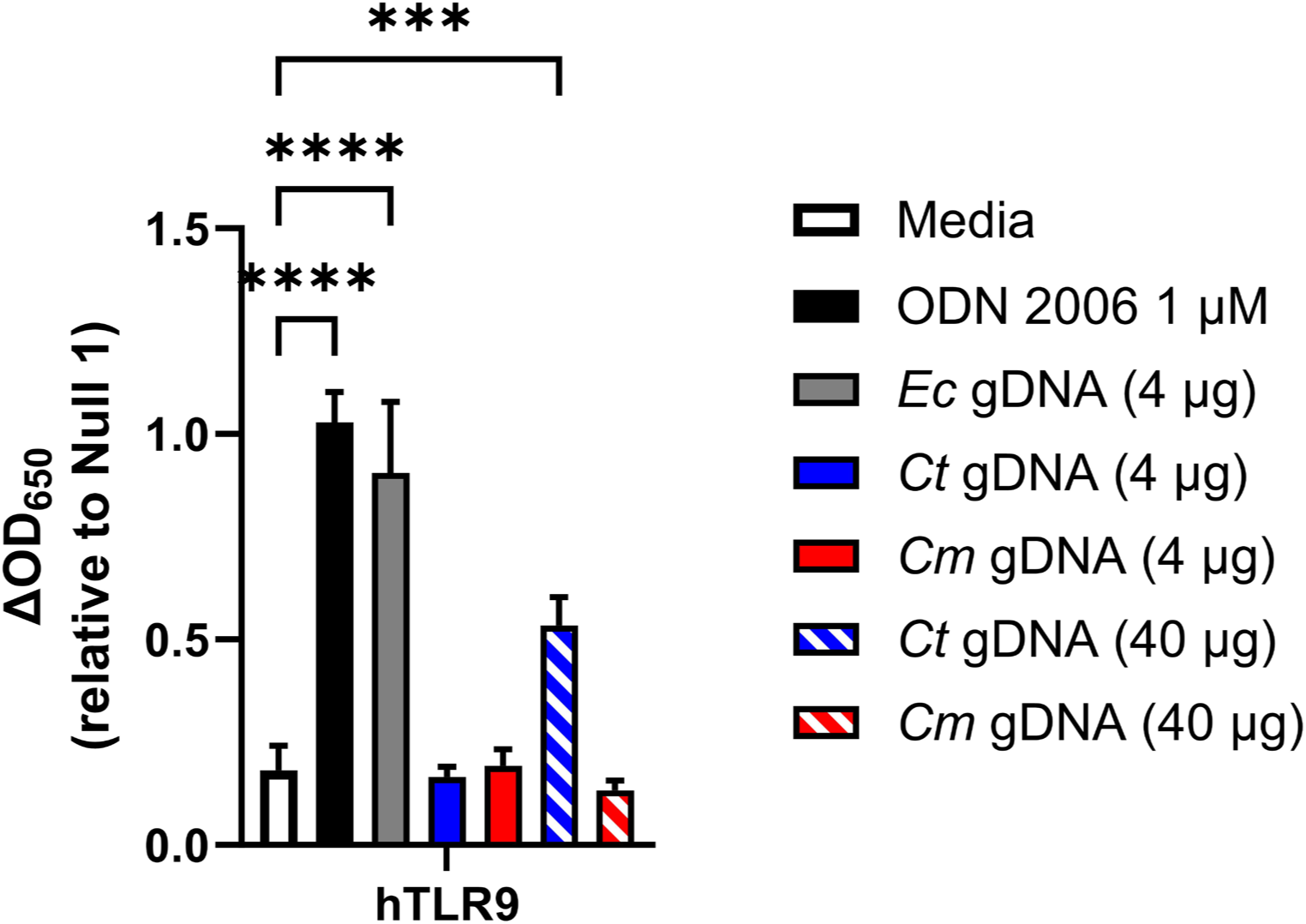
Genomic DNA from *C. trachomatis* induces hTLR9 signaling in vitro. A human TLR9 (hTLR9) HEK 293 reporter system was used to evaluate the stimulatory potential of sonicated genomic DNA isolated from *Escherichia coli* (strain MG1655), *C. trachomatis* (strain L2 434/Bu), and *Chlamydia muridarum* (strain Nigg). Data presented are the mean of three independent, biological replicates and error bars represent standard error of the mean. Groups were compared via one-way ANOVA with multiple comparisons. ****, p ≤ 0.0001; ***, p ≤ 0.001. All comparisons to the media control not shown were not significant.

### hTLR9 co-localizes with C. trachomatis in U937 cells

In unstimulated immune cells, TLR9 is maintained in the endoplasmic reticulum^20,40^. Upon stimulation, the receptor traffics through the Golgi Complex and primarily localizes to endolysosomes^41^, where it can interact with any foreign bacterial DNA sampled from the environment. While chlamydial inclusions are generally thought to be non-fusogenic with endosomes and lysosomes in most cell types^21–27^, the exception to the rule is macrophages and certain specialized dendritic cells^28–31^. While TLR9 is thought to be expressed in only subsets of dendritic cells and B cells in humans^42^, previous studies have demonstrated CpG-oligodeoxynucleotides (ODNs) signaling in the human monocyte-derived cell line U937^43–45^. To assess whether Chlamydia resides within TLR9-containing compartments within U937 cells, we conducted an immunolabeling experiment with cells infected with *C. trachomatis*. At 40 hpi, hTLR9 colocalizes with *C. trachomatis* inclusions in U937 cells **(Fig. 2A)**, indicating that this receptor successfully traffics to the inclusion in this cell line. Infected HeLa cells were used as a negative immunolabeling control, and no hTLR9 labeling was observed in the proximity of inclusions or anywhere in the infected cells **(Fig. 2B)**. These observations indicate that both *C. trachomatis* and hTLR9 associate within the same intracellular vacuole within this human monocyte-derived cell line.

**Figure 2.**
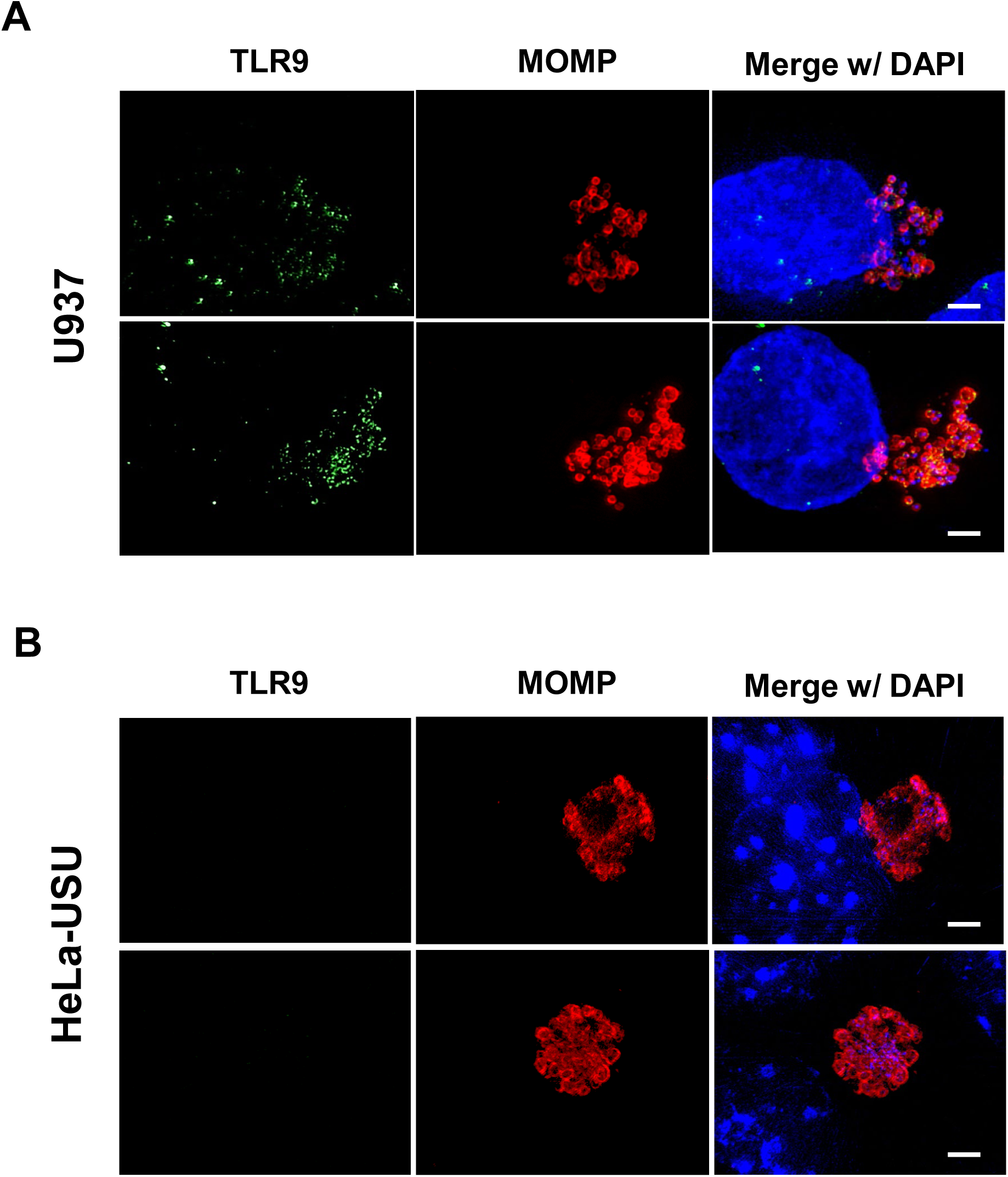
hTLR9 localizes to *C. trachomatis* inclusions in U937 cells. (**A**) U937 (pro-monocytic, human myeloid leukemia derived) and **(B)** HeLa cells were infected via rocking incubation with *C. trachomatis* at a MOI of ∼5. At 40 hpi, cells were fixed in 10% PFA, blocked with 3% BSA, and labeled with polyclonal antibodies to the pathogen’s <u>M</u>ajor <u>O</u>uter <u>M</u>embrane <u>P</u>rotein (MOMP) and TLR9. Images presented are Maximum Intensity Projections from zStacks acquired from a Zeiss PS.1 ELYRA imaging system and are representative of over 40 fields of view observed over two separate labeling experiments. Scale bar ∼5 µm.

### Pre-exposure to stimulatory TLR9 ligands has a minor but measurable effect on the development of C. trachomatis in U-937 cells

While *C. trachomatis* gDNA does stimulate signaling through hTLR9, it is significantly less immunostimulatory when compared to other bacteria **(Fig 1**, ^33^). Previous researchers have proposed that this represents a shared pathoadaption by bacterial STIs, enabling them to suppress their immunogenicity and evade or delay cell-mediated immune responses^33^. We sought to test this hypothesis by examining whether pre-treatment of U937 cells with a known hTLR9 agonist (ODN 2006) resulted in higher tolerance to subsequent infection by *C. trachomatis*. We found that cells pretreated with ODN 2006 appeared to be just as susceptible to infection as non-pretreated cells, as measured by inclusion forming unit counts at 24 hpi (**Fig. 3A**), however, there was a measurable difference (p<0.05) in the overall size of inclusions between the two groups, with fewer large (> 5µm diameter) inclusions present in the pretreated group (**Fig. 3B**). The vast majority of cells under both conditions contained small inclusions with only a few bacteria present, and the larger inclusions appeared oddly-shaped containing RBs that appeared dispersed and not closely associated with inclusion walls (**Fig. 3C**).

**Figure 3:**
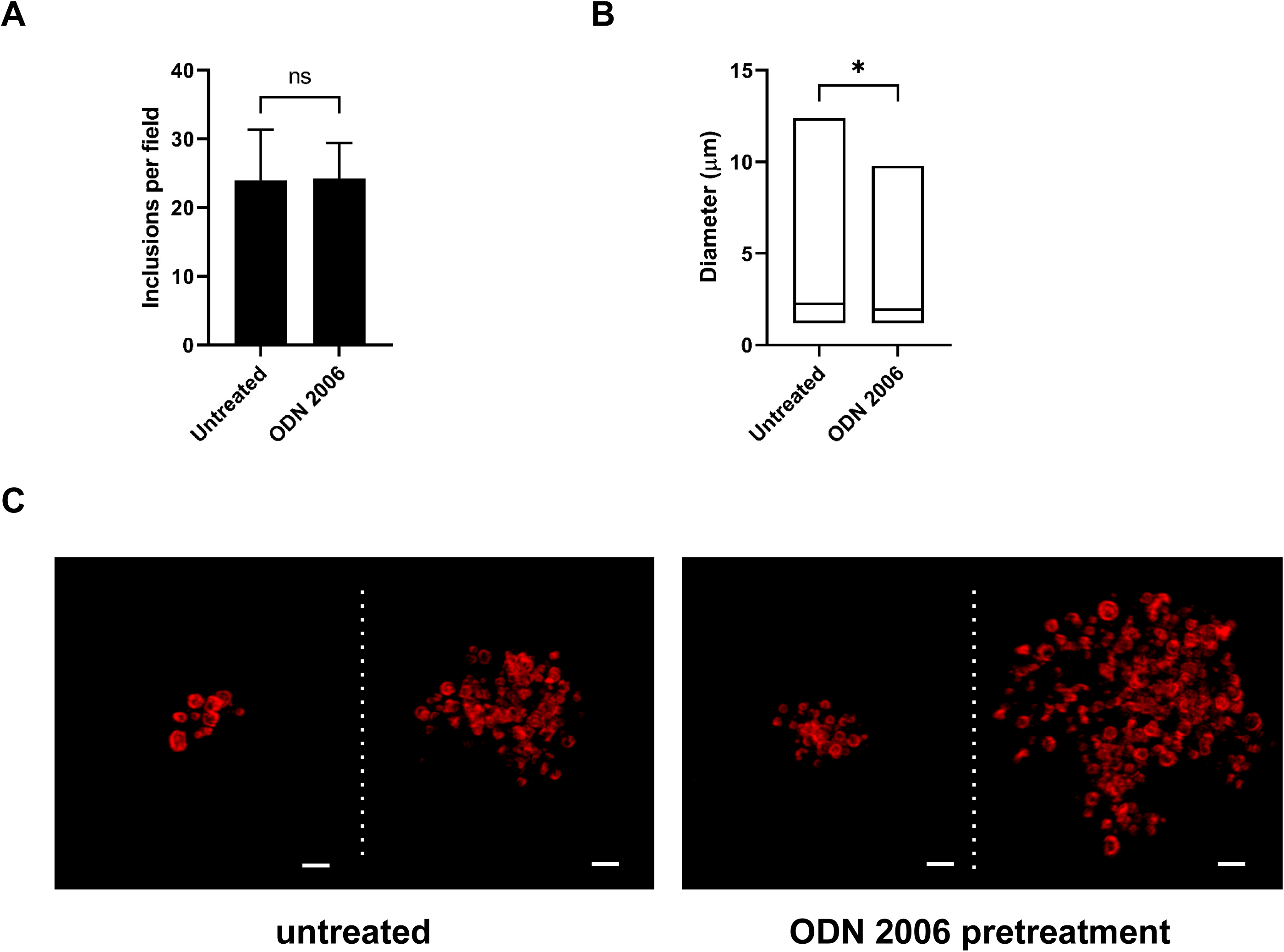
Pre-stimulation with a TLR9 agonist has a minor but measurable impact on the growth / development of *C. trachomatis* in the TLR9-expressing cell line U937. (**A**) The average number of inclusions observed per field of view (from 20 fields) in U937 cells that were left untreated or pretreated with the TLR9 agonist ODN 2006 for 18 hours prior to infection with *C. trachomatis*. Cell monolayers were fixed and labeled at 24 hpi. **(B)** Inclusion size measurements in U937 cells that were untreated or pretreated with ODN 2006 prior to infection. Data from two separate experiments were pooled for the analysis, and groups were compared via unpaired t test with Welch’s correction. *, p< 0.05. **(C)** Representative images of larger (≥ 5 µm) inclusions containing multiple RB-sized bacteria found in *Chlamydia*-infected U937 cells. All cells were fixed and imaged 24 hpi. Scale bar, ∼2 µm.

### Infection with live *C. trachomatis*, but not live *C. muridarum*, induces hTLR9 signaling in HEK293 reporter cells late in the pathogen’s developmental cycle

While we effectively demonstrated that chlamydial gDNA is immunostimulatory and recognized by TLR9 (**Fig. 1**), we reasoned that because the trafficking of extracellular gDNA to TLR9-containing vesicles is dependent on endosomes^19^, this would likely be the case for Chlamydia-specific TLR9 signaling as well. However, as chlamydial inclusions only associate with endosomes / lysosomes in dendritic cells, and our hTLR9 reporter cell line is not of monocytic origin^46^, we questioned whether direct infection of hTLR9-expressing HEK-293 cells with *C. trachomatis* would result in measurable, hTLR9-specific SEAP activity. To test this, reporter cells were infected with *C. trachomatis* at a MOI of either 2 or 0.2 and SEAP activity was assessed at 24 and 40 hours post infection (hpi). Surprisingly, direct infections did result in measurable, dose-dependent hTLR9 activity, albeit significant signaling did not occur until the later stages of the pathogen’s developmental cycle (**Fig. 4A**). Follow-up assays demonstrated that this signaling was unique to *C. trachomatis*, as live infections with *C. muridarum* did not result in similar hTLR9-specific activity (**Fig. 4B**). To account for the possibility that *C. trachomatis*-specific hTLR9 signaling was due to the presence of extracellular gDNA from killed / lysed bacteria present in the infection inoculum, we ran the assay comparing the signaling potential of live vs. heat-killed organisms, and found measurable TLR9 signaling activity only during live infections (**Fig. 4C**). These observations indicate that chlamydial viability is required for this late-stage signaling observed in our hTLR9 reporter cells.

**Figure 4:**
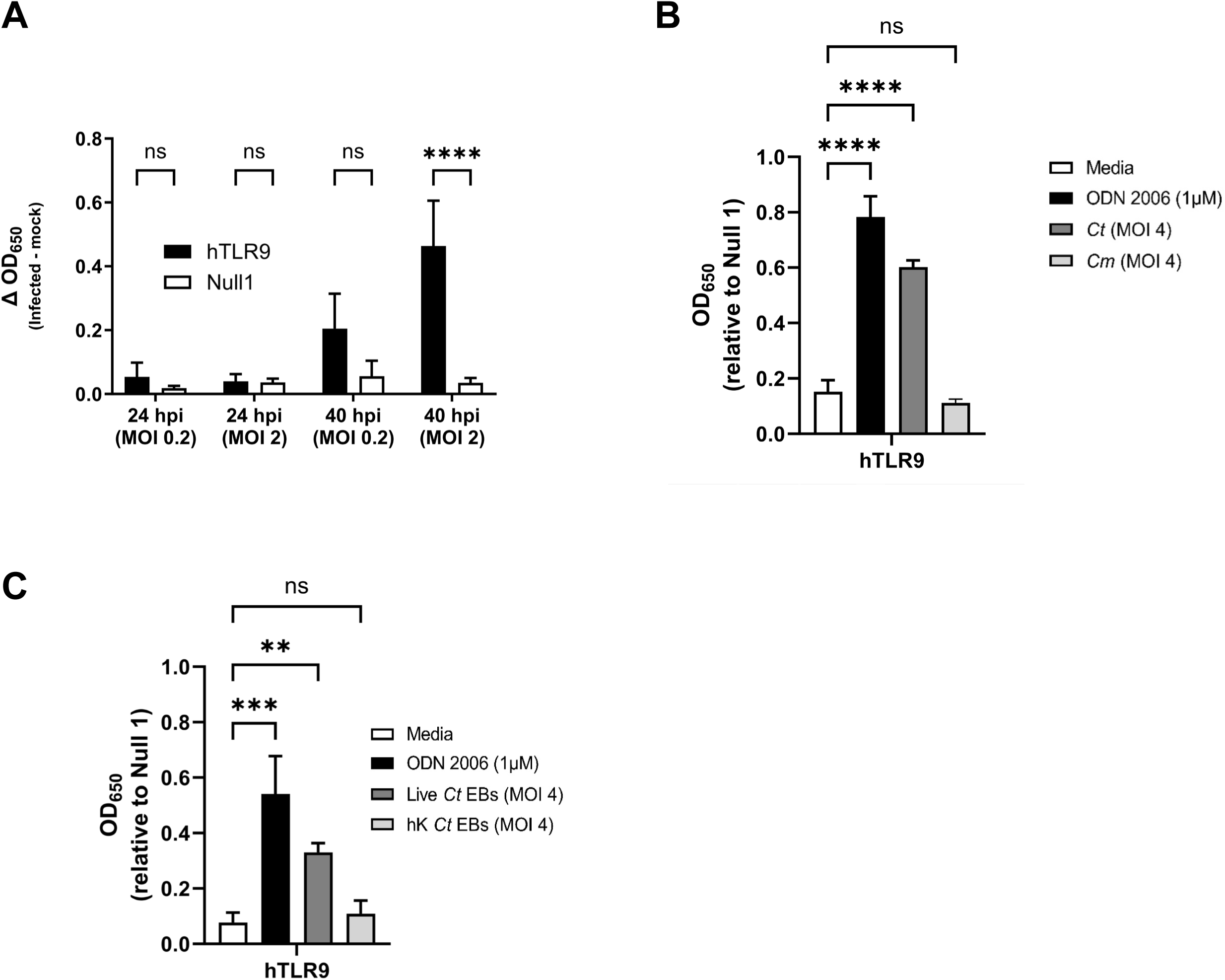
*C. trachomatis*-induced hTLR9 signaling occurs late in the pathogen’s developmental cycle in non-phagocytic cells. (**A**) SEAP activity was measured from the supernatants of hTLR9– and Null 1-HEK 293 reporter cells infected with *C. trachomatis* serovar L2 (strain Bu/434) at 24 and 40 hpi. **(B)** A comparison of the hTLR9-dependent SEAP activity present in supernatants obtained from reporter cells infected with either *C. trachomatis* or *C. muridarum* for 48 hours. **(C)** hTLR9 signaling assessed at 48 hpi for live and heat-killed *C. trachomatis* EBs. For all panels, columns represent the mean value calculated for data acquired from three separate experiments (biological replicates) and error bars represent standard error of the mean. Groups were compared via two-way and one-way ANOVA with multiple comparisons, respectively. ****, p ≤ 0.0001; ***, p ≤ 0.001; ns, not significant.

### Both live *C. trachomatis* and *C. muridarum* signal through mTLR9 signaling in HEK293 reporter cells

We found it curious that *C. trachomatis* appeared to signal more robustly through hTLR9 than *C. muridarum*, as we had hypothesized that each pathogen would have evolved to minimize the immunogenicity to the TLRs of their respective host species. To examine this question of cross-species recognition further, we conducted an additional strain comparison using a reporter cell line expressing the murine allele of TLR9 (mTLR9). mTLR9 has been shown to exhibit structural differences from hTLR9 and recognizes different CpG motifs^42,47^. We examined three different MOIs for each organism (10, 1, and 0.1), and observed mTLR9 signaling at the 24 hpi time point for both *C. trachomatis* or *C. muridarum* when our highest MOI was used **(Fig. 5A)**. At the 48 hpi time point, mTLR9 signaling was observable at MOIs as low as 1 (**Fig. 5B**), and when compared directly, *C. trachomatis* signaling was approximately twice that of *C. muridarum* at both time points examined (**Fig. 5C, D**). When combined with the purified gDNA data, these observations demonstrate that *C. muridarum* is significantly less stimulatory than *C. trachomatis* as assessed by both the human and murine TLR9 receptors.

**Figure 5:**
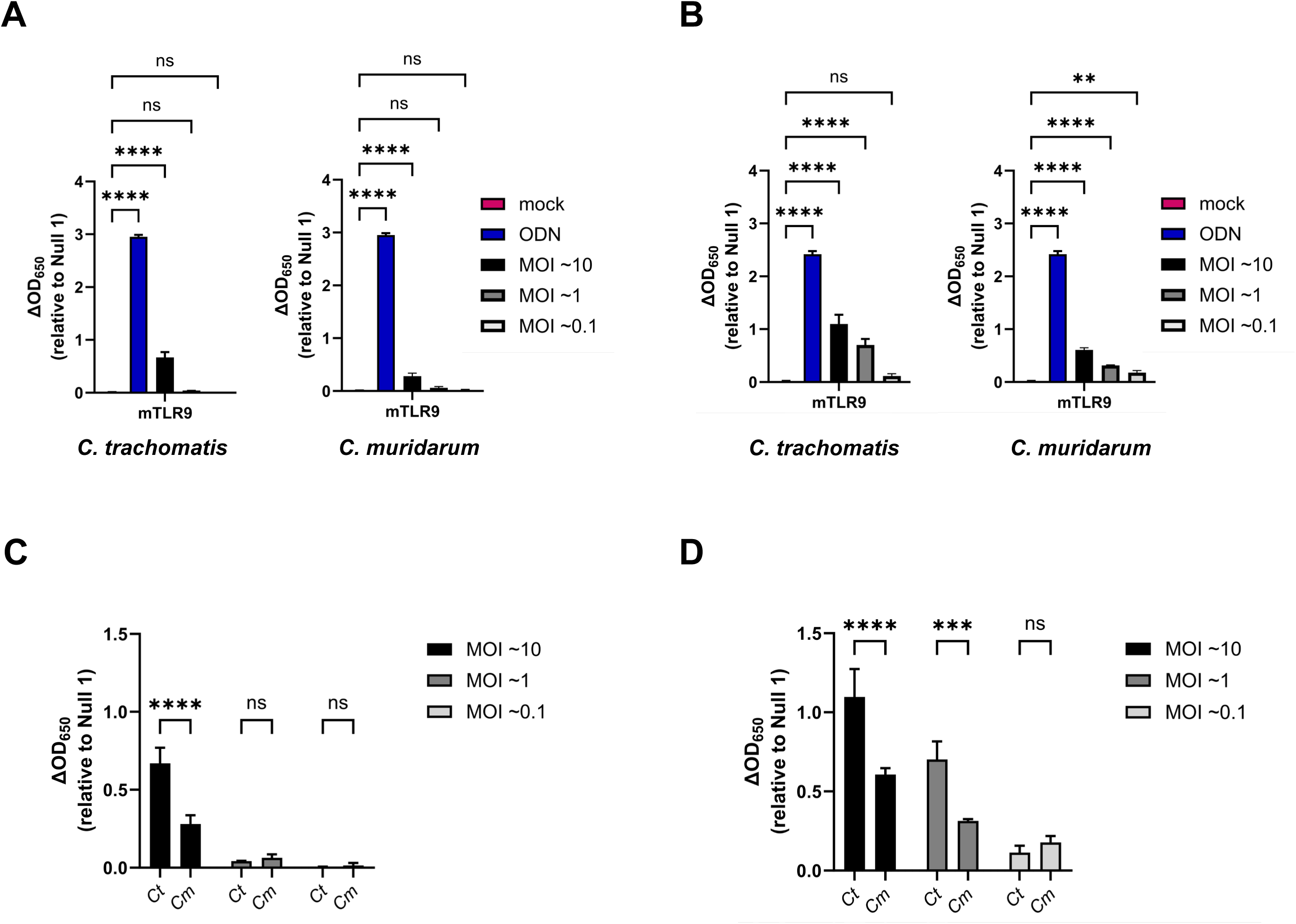
Both *C. trachomatis* and *C. muridarum* signal through mTLR9 during live infections of HEK293 reporter cells. SEAP activity was measured from the supernatants of mTLR9-HEK 293 reporter cells infected with *C. trachomatis* and *C. muridarum* at 24 **(A)** and 48 hpi **(B)**. A side-by-side comparison of the stimulatory activity of each strain is presented for 24 **(C)** and 48 **(D)** hpi. For all panels, columns represent the mean value calculated for data acquired from three biological replicates and error bars represent standard error of the mean. Groups were compared via one-way and two-way ANOVA with multiple comparisons, respectively. ****, p ≤ 0.0001; ***, p ≤ 0.001; **, p<0.01; ns, not significant.

### Chlamydia-specific TLR9 signaling in HEK293 cells occurs as a result of the RB-to-EB conversion

Given that we did not observe Chlamydia-specific hTLR9 signaling at 24 hpi **(Fig. 4A)**, we reasoned that this could be due to one of two possibilities: 1) that not enough intracellular bacterial replication had occurred to meet the threshold needed for detection by TLR9 and / or 2) that chlamydial gDNA availability varied at different time points post infection. The conversion between the chlamydial replicative and infectious forms begins to occur (in cultured epithelial cells) between ∼20-24 hpi^48,49^. To determine whether the interruption of chlamydial EB development affects Chlamydia-specific TLR9 signaling, we measured signaling in Chlamydia-infected cells that were treated with chloramphenicol (Cm_25_) at various time points post-infection. We observed that signaling only occurred in untreated cells and cells treated with Cm_25_ at or later than 22 hpi (**Fig. 6A**), roughly coinciding with the initiation of RB-to-EB conversion.

**Figure 6.**
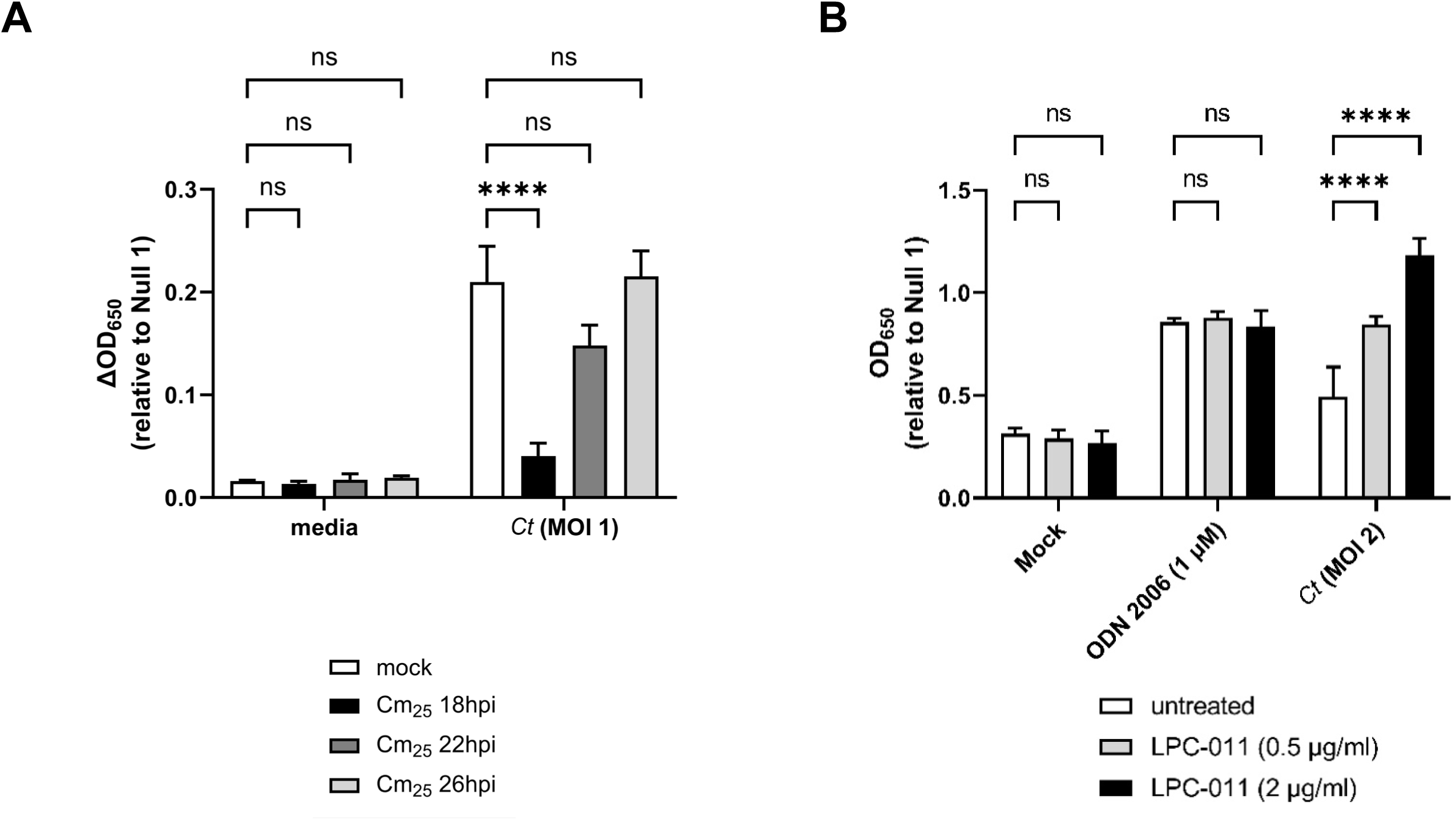
Chlamydia-specific TLR9 signaling in HEK293 cells occurs as a result of the RB-to-EB conversion. (**A**) *C. trachomatis*-infected hTLR9-HEK293 reporter cells were treated with chloramphenicol (25 ug/mL; Cm_25_) at the time points indicated and SEAP activity was measured at 44hpi. **(B)** The effects of the LOS inhibitor LPC-011 on *C. trachomatis*-induced hTLR9 signaling. All columns represent mean values calculated for data acquired from three biological replicates and error bars represent standard error of the mean. Groups were compared via two-way ANOVA with multiple comparisons. ****, p ≤ 0.0001; ns, not significant.

The autolysis of *C. trachomatis* RBs has been described previously^50^, and it is commonly thought that a small subset of RBs within the inclusion will undergo lysis during the developmental cycle. We hypothesized that such lytic events are likely the cause of gDNA release within inclusions. Given the delay we observed in TLR9 signaling in our HEK293 report cells (**Fig. 4A**), we also hypothesized that the most likely time when gDNA is released from the microbe would be during the transition state between replicative and infectious forms. To examine the degree to which gDNA release impacted our Chlamydia-specific TLR9 signaling, we examined the effects of the LpxC inhibitor LPC-011^51^ on *C. trachomatis*-induced TLR9 signaling. LpxC carries out the 2^nd^ step in the lipooligosaccharide (LOS) biosynthesis pathway, and while LOS biosynthesis is not required for the transition of Chlamydia EBs to RBs, nor for RB replication, it is required for the conversion of RBs into EBs at the end of the pathogen’s developmental cycle^51,52^. In the absence of LOS, RBs lyse during the conversion process. When infected HEK293 cells were treated with the LOS inhibitor LPC-011, a dose-respondent increase in Chlamydia-specific hTLR9 signaling was detected (**Fig. 6B**). Together, these observations support the hypothesis that *C. trachomatis* gDNA release in epithelial cells is the result of the process of RB-to-EB conversion.

### Persistence alters Chlamydia-induced hTLR9 signaling

We have previously reported that stress-inducing conditions linked to the aberrant / persistent phenotype in *C. trachomatis* can directly affect the recognition of the pathogen by the Nod-like Receptor (NLR) NOD1^53^. Prior studies have also found that these persistence-inducing conditions can affect the ability of Chlamydia and Chlamydia-related organisms to regulate DNA replication. While cell division is inhibited during persistence, some persistence-inducing conditions result in a marked reduction in chlamydial DNA replication^54,55^ while others result in the accumulation of 100s of genome copies residing within a single, enlarged bacterium^55–59^. Given that many of these ‘aberrant’ chlamydial forms exhibit evidence of membrane disruptions and structural defects^60–62^, we reasoned that DNA could likely be shed by these aberrant forms into the inclusion space. We also hypothesized that continued DNA replication would directly impact TLR9-stimulation by *C. trachomatis*, as the abundance of available ligand (gDNA) would likely correspond to the signaling potential of the microbe. We began by examining genome replication in *C. trachomatis* under aberrance induction by ampicillin and the iron chelator 2,2′-Dipyridyl (Dpd) over the course of the microbe’s developmental cycle (∼44hpi). Similar to previously published results^48^, we found that the majority of *C. trachomatis* genome replication occurs within the first 24 hours post-infection under all the conditions tested, and that persistence induced by iron chelation resulted in ∼1 log_10_ reduction in genome counts **(Fig. 7A)**.

**Figure 7.**
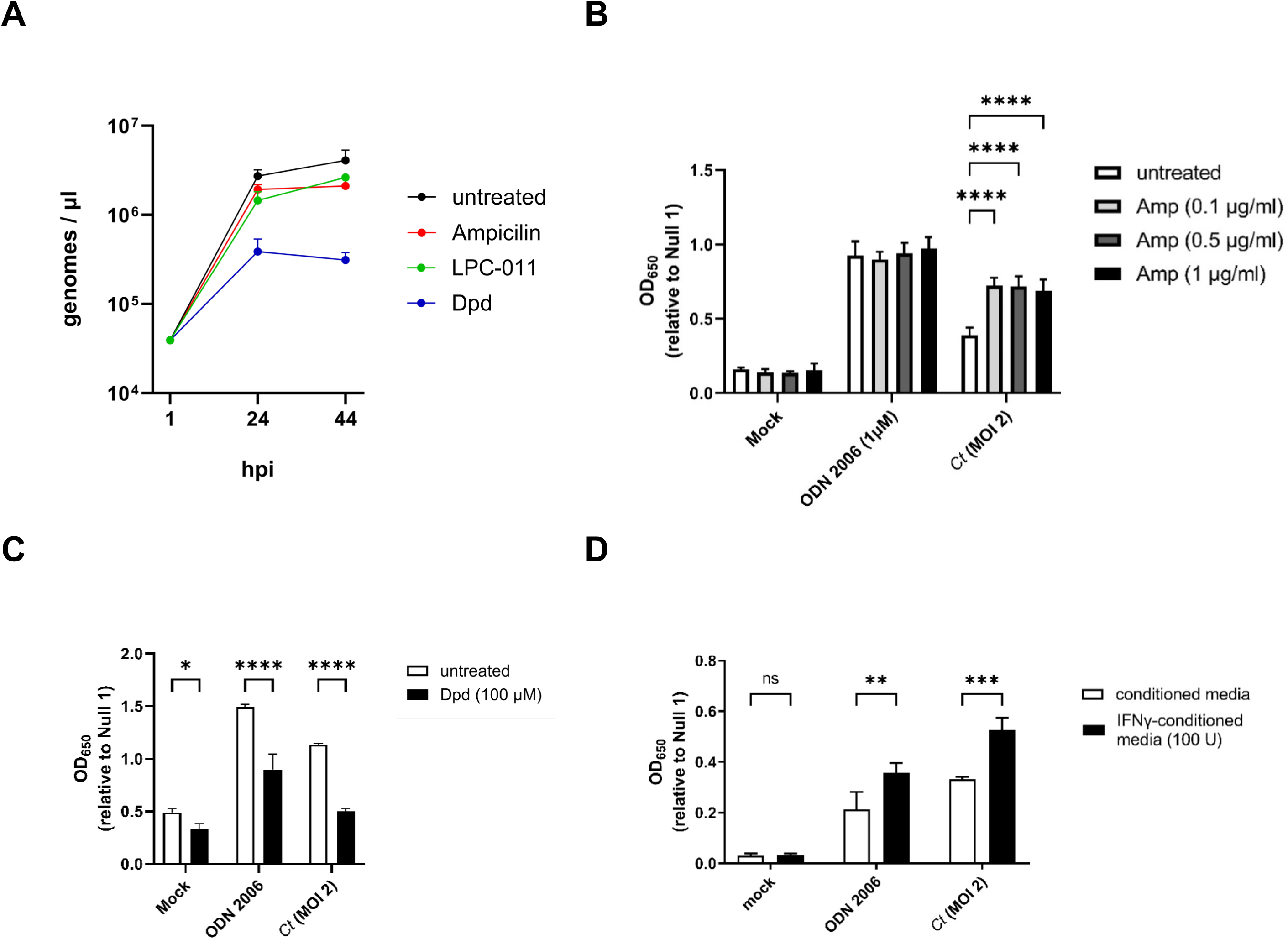
Persistence alters Chlamydia-induced hTLR9 signaling. (**A**) *C. trachomatis* genome copies were measured in untreated cells as well as in cells infected in the presence of ampicillin, the iron chelator 2,2′-Dipyridyl (Dpd), and the LpxC inhibitor (LPC-011) over the span of 44 hours. Measurements were taken at 1, 24, and 44 hpi. **(B-D)** The effects of **(B)** ampicillin; Amp, **(C)** 2,2′-Dipyridyl; Dpd, and **(D)** tryptophan-depletion via interferon gamma (IFNγ) were accessed on hTLR9-stimulatory activity of ODN 2006 and infection by *C. trachomatis*. All columns represent mean values calculated for data acquired from three biological replicates and error bars represent standard error of the mean. Groups were compared via two-way ANOVA with multiple comparisons. ****, p ≤ 0.0001; ***, p ≤ 0.001; **, p < 0.01; *, p ≤ 0.05; ns, not significant.

Having assessed the relative differences in genome replication under a variety of persistence-inducing conditions, we next tested Chlamydia-infected cells under these same conditions over a 44-hour period in our hTLR9 HEK293 reporter assay. Antibiotics that target peptidoglycan are known to result in high levels of ‘temporary polyploidy’ in the resulting chlamydial aberrant bodies^55–59^ and we observed that in the presence of ampicillin, *C. trachomatis*-induced TLR9 activity was significantly enhanced **(Fig. 7B).**

Conversely, iron restriction is associated with a reduction in genome replication in *Chlamydia* and Chlamydia-like organisms (**Fig. 7A**,^54,55^) and we observed a marked decrease in TLR9 activity when infected cells were treated with the iron chelator 2,2-dipyridyl (Dpd) **(Fig. 7C)**. However, we also observed that the condition impacted our positive control (ODN 2006), indicating that the effects on signaling are likely indirect. Tryptophan starvation resulting from treatment of primate-derived cells with interferon gamma (IFNγ) also results in the accumulation of large numbers of genome copies within single aberrant chlamydial bodies^63,64^, and we found that this condition significantly enhanced hTLR9 signaling **(Fig. 7D)**, however, this was also not unique to *Chlamydia*-induced signaling as we saw significant enhancement in our positive control. Taken together, these data indicate that TLR9 signaling results from the release of gDNA from *C. trachomatis* within its inclusion and that, similar to NOD1 signaling^53^, ‘persistence’-inducing conditions can affect both Chlamydia-specific and Chlamydia non-specific TLR9 signaling.

## DISCUSSION

Our data indicates that *C. trachomatis* gDNA is a stimulatory ligand for both human and murine TLR9 and has a higher stimulatory potential than the murine pathogen *C. muridarum*. We found that TLR9 signaling from *C. trachomatis* was higher than that of *C. muridarum* in both *in vitro* testing of purified gDNA **(Fig. 1)** and live infections of reporter cells **(Fig. 4B and Fig. 5)**. Given our data indicating that TLR9 signaling is impacted by the timing of the chlamydial developmental cycle **(Fig. 4A and Fig. 6A)** and bacterial load **(Fig. 4A and Fig. 5)**, this is a particularly significant finding. *C. muridarum* has a higher replication rate and a shorter developmental cycle than *C. trachomatis*^65^, and thus likely releases gDNA into the inclusion earlier and at a higher abundance than *C. trachomatis*. The fact that *C. muridarum* TLR9 signaling is so much lower than *C. trachomatis* is likely the result of fewer CpG-stimulatory motifs present in its genome than are present in *C. trachomatis*. This difference in the immunostimulatory potential of the gDNA from of these two pathogens may have arisen due to differences in tissue/host-specific prevalence of TLR9. TLR9 was originally thought to be expressed in a number of cell types (such as macrophages) in mice^66^ and expressed in only a few specialized dendritic cell populations in humans^67^. However, more recent studies have brought this general consensus into question, and TLR9 is now known to be highly expressed in multiple cell types in both human and murine lung tissue^68^. Given its origin^69^ (and re-emergence^70^) as a murine lung pathogen, *C. muridarum* likely evolved under heightened selective pressure in this environment and the reduced stimulatory potential of its gDNA may be evidence of a pathoadaptation to this niche.

TLR9 haplotypes and polymorphisms have been previously shown to correlate with potential risk of cervicitis^35^, cervical cancer^71^, as well as susceptibility and the development of symptoms after HR-HPV and *C. trachomatis* infection^36,72^. TLR9-stimulatory adjuvants have also been used extensively in chlamydial vaccine development^73–77^. Despite this, TLR9 has been deemed to play only a minor role in chlamydial recognition during active infections. This general consensus has been based on studies investigating pattern recognition molecules generated by *C. muridarum* that are recognized by oviduct epithelial cell lines^38^ and assessing the impact of murine TLR9 on the clearance of *C. trachomatis* and *C. pneumoniae* infections utilizing a TLR9−/− mouse infection model^34,39^. While these studies were well powered and well-controlled, they did not directly explore the interaction of either pathogen with the receptor at early stages of infection. The relatively high doses required to generate stable *C. trachomatis* infections in the murine model are likely due to the presence of innate immunological factors (such as TLR9) that are thought to play a larger role in preventing infections from occurring (particularly in the lower genital tract), rather than controlling them once they are established. By comparison, TLR2^38,78–82^ and TLR4^7–9,83^ are thought to play a critical role in controlling chlamydial infections. While TLR2 and TLR4 have been shown to be highly expressed in the upper genital tract, TLR9 as well as NOD-like Receptors (NLRs) 1 and 2 are expressed more uniformly throughout the upper and lower genital tract^84^. Similarly, the cGAS-STING signaling pathway, another innate DNA recognition system that grants heightened immunity to *C. trachomatis* infection, is notably restricted to the lower genital tract^85^.

Given that we observed co-localization of *C. trachomatis* inclusions with TLR9 in U937 cells **(Fig.2)**, the availability of chlamydial gDNA in this circumstance is understandable. In macrophages and dendritic cells, the AP3 adaptor complex and the AP-3-interacting cation transporter (Slc15a4) are responsible for trafficking TLR9 from the early endosome to a specialized lysosome-related organelle where TLR9 then activates MYD88 signaling^86^. In contrast, while a number of TLRs (as well as their adaptor protein MYD88) associate with the chlamydial inclusions in epithelial cells^78,87^, these inclusions avoid fusing with endosomes / lysosomes^88^. Thus, while the mechanism for TLR9 interacting with gDNA from *Chlamydia species* in U937 cells is straightforward (bacterial lysis in lysosomes containing TLR9), the mechanism by which signaling occurs during live infections in our HEK293 report cells is not. Given that we observe signaling only on the second day post infection **(Fig. 4A)**, we believe that TLR9 signaling under these conditions is the result of gDNA being released from a subset of chlamydial RBs that fail to successfully convert into EBs. Our data demonstrating 1) prevention of the RB to EB conversion eliminates TLR9 signaling **(Fig. 6A)** and 2) artificially enhancing the frequency of lysis events during conversion (utilizing the LpxC inhibitor) enhances TLR9 signaling **(Fig. 6B)** supports this proposition.

It is presently unclear how the gDNA that is released during RB lysis traffics to TLR9-containing endosomes in our HEK293 cells. Previous studies have indirectly demonstrated the presence of cytosolic chlamydial gDNA in infected cells in culture^89,90^, and we hypothesize that if chlamydia gDNA is capable of exiting the inclusion into the cell cytosol, this might enable it to eventually be trafficked to TLR9-containing compartments. To date, no direct mechanism has been proposed by which chlamydial DNA exits the inclusion. Evidence exists in multiple model systems that DNA transfer from a bacterium to a host cell can occur via type IV secretion systems (T4SS)^91–93^, but there exists no example of DNA being successfully transferred via the secretion systems utilized by *C. trachomatis* (ie. type II and type III secretion). Experiments investigating alternative trafficking route(s) of chlamydial gDNA from the inclusion to the TLR9 receptor and currently ongoing.

In addition to immune recognition, the presence of cytosolic chlamydial DNA in infected host cells has implications for other aspects of chlamydial biology, such as factors that influence genetic exchange between strains^94^ and potential barriers to inter-species gene flow. *Chlamydia* species undergo conversion from RBs to EBs in a rather disjointed fashion, and the reasons for this are not well understood. *C. trachomatis* encodes ComEC homologs that enable DNA uptake and lateral gene transfer^95^. These are presumably expressed predominantly in RBs, as most metabolic processes in chlamydial EBs are significantly reduced. Given our observations that *C. trachomatis* releases gDNA into its immediate environment during RB-to-EB transition events **(Fig 6)**, it stands to reason that asynchronous conversion between developmental forms may have arisen, in part, due to the potential for genetic exchange between replicating RBs and RBs lysing during conversion. *C. trachomatis* inclusions exhibit homotypic fusion, in that multiple inclusions within a cell can fuse into a single, enlarged inclusion. This process is largely dependent on IncA^96–98^, a SNARE-like protein that is secreted by the chlamydial Type III secretion system and incorporates into the inclusion wall. Infections with different strains^99,100^ and mixed infections with different chlamydial species^101,102^ have been used to date to demonstrate the potential gene flow between Chlamydia species that are capable of residing within shared inclusions. Interestingly, genetic exchange between non-fusogenic species has also been observed^102^, suggesting that DNA trafficking between inclusions can also occur. Studies exploring the mechanism(s) by which gDNA transits to and from the host cell cytosol are presently ongoing.

## Conclusions

In this study, we demonstrate the stimulatory potential of Chlamydial gDNA for recognition by hTLR9 and mTLR9, show that hTLR9 colocalizes with chlamydial inclusions in the pro-monocytic cell line U937, demonstrate that under normal growth conditions Chlamydia-specific TLR9 signaling occurs during the stage in the pathogen’s developmental cycle when developmental form conversion occurs, and that the induction of aberrance can either enhance or diminish this signaling. The disjoining of genome replication and cell division that occurs during chlamydial persistence has been previously suggested to be a beneficial adaptation, enabling the microbe to rapidly proliferate via numerous, simultaneous cell division events upon the removal of stress-inducing conditions. Our data indicates that any such benefit likely comes with inherent costs in immune cells, specifically the recognition of immunostimulatory chlamydial gDNA by a host’s innate immune defenses. We reason that persistence requires the evasion of immunological clearance mechanisms and as such, models of chlamydial persistence (to include those independent of the ‘aberrant’ phenotype^103^) that demonstrate this characteristic in vitro are more likely to correspond to phenotypes associated with persistence in animal models and human patients.

## MATERIALS AND METHODS

### Reagents

Anti-TLR9 (AB134368) was purchased from Abcam and Invitrogen, respectively. Anti-MOMP (LS-C123239) was purchased from LSBio. LpxC inhibitor LPC-011 was graciously provided by Dr. Pei Zhou (Duke University).

### Bacterial Strains and Cell Lines

*C. trachomatis* serovar L2 strain 434/Bu, *C. muridarum* strain Nigg, and *E. coli* strain MG1655 were provided by Anthony Maurelli (University of Florida). Chlamydial stocks were generated utilizing HeLa-USU cells (also provided by Anthony Maurelli) unless otherwise noted. Whole cell lysate (‘crude’) freezer stocks of chlamydial EBs were generated from HeLa cells 40 hours post infection and stored at –80° C in sucrose phosphate glutamic acid buffer (7.5% w/v sucrose, 17 mM Na_2_HPO_4_, 3 mM NaH_2_PO_4_, 5 mM L-glutamic acid, pH 7.4) until use. Stocks were titered via inclusion forming unit (IFU) assay (described below). HEK-Blue-hTLR9, –mTLR9, and –Null1 cells were purchased from InvivoGen and propagated according to the manufacturer’s instructions. Cell lines were passaged in high-glucose Dulbecco’s modified Eagle medium (DMEM; Gibco) and 10% fetal bovine serum (FBS; HyClone). U937 cells were purchased from ATCC and propagated according to their instructions. All cell lines were checked for mycoplasma contamination 2 passages after the initial liquid nitrogen thaw, and every subsequent 10 passages.

### Genomic DNA isolation

Prior to the day of isolation, one 7 ml overnight culture of *E. coli* strain MG1655 was prepared. On the day of isolation, (4) 100 µl aliquot ‘crude’ EB preparations of *C. trachomatis* serovar L2 strain 434/Bu and *C. muridarum* strain Nigg were thawed at room temperature. As stocks thawed, 6 ml of the overnight *E. coli* culture was centrifuged at 16,000 G for 2 minutes. gDNA from *C. trachomatis*, *C. muridarum*, and *E. coli* were then isolated using a Promega Genomic DNA Purification Kit. gDNA from each species was rehydrated in 50 µl of sterile, deionized water (Fischer) and placed in a 4° C fridge overnight. On the following day, aliquots were placed on ice and the concentration of gDNA was assessed via nanodrop. To sufficiently shear gDNA for use in TLR9 signaling assays, gDNA stocks were then pulse-sonicated ∼10 times (sonicating for 3 seconds then pausing for 1 – 2 seconds), and placed on ice immediately after for at least 10 minutes. If not used the same day, gDNA stocks were then stored at –20° C.

### Quantification of Inclusion Forming Units (IFU)

96 well tissue culture-treated plates (Fisher) are seeded 24 hours prior to IFU assays with 200µl per well of a 200,000 L2 cells/ml suspension in DMEM / 10% FBS (∼40,000 cells per well). On the day of the assay, bacterial suspensions are thawed on ice and then serially diluted in infection medium [DMEM, 10% FBS, MEM Non-Essential Amino Acids (Sigma), 0.5µg/ml cycloheximide]. Spent media is removed from the 96 well plate and 200 µl of each chlamydia dilution is added to each well, with each dilution conducted in duplicate.

Plates were then spun in a tabletop centrifuge (Eppendorf) at 3000 rpm at 35° C for 1 hour to synchronize infection and ensure that all infectious EBs come into contact with cell monolayers. Plates were then incubated for 24-28 hours at 37° C 5% CO_2_. At the desired hpi, infection medium was removed by suction, infected cells were fixed / permeabilized by the addition of 200 µl of ice-cold methanol and incubated at room temperature for ∼10 minutes. The methanol was then aspirated and 1 drop of Pathfinder Chlamydia Culture Confirmation System (BioRad) was added to each well. Plates were incubated at room temperature in the dark for 30 minutes, after which time the antibody staining solution was removed, wells were gently washed 3 times with deionized water, and 1 drop of glycerol mounting medium was added to each well. Plates were inverted, stored at 4° C and examined under an epifluorescence microscope (Olympus) within 24 hours. Total inclusion forming units (IFU) were calculated by counting the number of visible inclusions for 20 different imaging fields using the 40x objective.

### *C. trachomatis* Infections of U937 cells

For infecting U937 cells, ∼2.5 x 10^5^ cells / mL were spun down and resuspended in infection medium containing ∼1×10^6^ IFU of *C. trachomatis* (MOI ∼4). 2.5 x 10^5^ cells were then added to each well of a 24 well plate and placed in the incubator for 24 or 44 hours. For pre-stimulation experiments, ∼2.5 x 10^5^ cells / mL were spun down and resuspended in infection medium + / – 1µM ODN 2006 and incubated overnight at 37° C. The next morning, cells were spun down and resuspended in infection medium containing ∼1×10^6^ IFU of *C. trachomatis* (MOI ∼4), 2.5 x 10^5^ cells were then added to each well of a 24 well plate and placed in the incubator (at 37° C) for an additional 24 or 44 hours.

### Fluorescence Microscopy / TLR9 and MOMP-labeling

Briefly, HeLa cells were infected with *C. trachomatis* L2 434/Bu as described above. At indicated time points infection medium was removed, and cells were washed three times with 1× PBS. Cells were fixed and permeabilized with methanol for 5 min and gently washed three times with 1× PBS. Cells were then further permeabilized and blocked as described above. Cells were then washed with 3% BSA and incubated with donkey anti-mouse or donkey anti-rabbit Alexa Fluor 488 (1:1,000 in 3% BSA). For visualizing cell nuclei, cells were incubated with Hoechst stain for ∼5 min and then washed with 3% BSA and 1× PBS.

For labeling TLR9 in infected U937 and HeLa cells, cells were fixed with 10% paraformaldehyde (PFA) for 10 min at room temperature and subsequently washed with 1× PBS. Cells were then permeabilized with 100% Methanol for 2 minutes, washed with 1× PBS, and then blocked for 1 h with 1% BSA. Cells were then incubated with anti-MOMP and anti-TLR9 (1:100) for 1 hour at room temperature, followed by several washes 3% BSA, and subsequent incubation with anti-goat and anti-rabbit conjugated antibodies (Alexa594 and Alexa488, respectively). Cells were then further washed with 3% BSA, PBS, and then coverslips were mounted on slides with ProLong gold antifade mounting medium and stored in the dark at 4°C prior to imaging via structured-illumination (Elyra PS.1) microscopy.

### HEK-Blue hTLR9 and Null1 NF-κB reporter assay

HEK-Blue cells expressing human or murine TLR9 and carrying the NF-κB SEAP (secreted embryonic alkaline phosphatase) reporter gene (InvivoGen) were used according to the manufacturer’s instructions and adapted to assess TLR-specific NF-κB activity induced via live *C. trachomatis* and isolated bacterial gDNA. Briefly, 3 × 10^5^ cells/ml of HEK-Blue-hTLR9 or –Null1 cells were plated in 96-well plates (total reaction volume of 200 μl per well [∼6.0 × 10^4^ cells per well]) and allowed to settle/adhere overnight at 37°C. Media was then removed and replaced with 200 µl of medium containing either *C. trachomatis* (at the MOIs indicated), purified genomic DNA, or the known TLR9-stimulating ligand ODN 2006. Plates were then centrifuged for 1 h at 2,000 × *g* and subsequently incubated in a CO_2_ incubator at 37°C. Cell supernatants were collected at indicated time points for subsequent analysis of SEAP activity. A colorimetric reporter assay was then utilized to quantify the abundance of SEAP in cell supernatants. Twenty microliters of supernatant collected from infected cells was added to 180 μl of the SEAP detection solution (InvivoGen), followed by incubation at 37°C for ∼6 h. SEAP enzymatic activity was then quantified using a plate reader set to 650 nm. Infected cells were compared to uninfected (media) controls and cells treated with ODN 2006 (positive control). To ensure that changes in alkaline phosphatase activity were TLR-dependent under each of the experimental conditions tested, all experiments were carried out in parallel in the HEK-Blue-Null1 cell line, which contains the empty expression vector but lacks TLR9.

HEK-Blue TLR9 SEAP reporter assays were always carried out in three separate experiments, statistical analysis was conducted by either 1– or 2-way ANOVA, and significance values were analyzed by utilizing Sidak’s multiple-comparison test.

### Assessing Chlamydial genome copy number

L2 cells in 24-well plates were infected with *C. trachomatis* L2 at MOI of 0.1 as described above. Total content of the wells was harvested at 1, 24 and 44 hpi and DNA was extracted using DNAeasy blood and tissue kit (Qiagen). Genome copy number quantitation was done by quantitative polymerase chain reaction (qPCR) using *C. trachomatis* 16S rRNA primer-probe mix (Forward 5’-GTAGCGGTGAAATGCGTAGA-3’, Reverse 5’-CGCCTTAGCGTCAGGTATAAA-3’, Probe 5’-ATGTGGAAGAACACCAGTGGCGAA-3’) (Integrated DNA Technologies), iTaq universal probes supermix (Bio-Rad) and 1 µl DNA as template. PCR reaction (40 cycles) was done on a CFX96 qPCR machine (Bio-Rad) using following conditions: initial denaturation 95°C for 3 min, denaturation 95°C for 5s and annealing/extension 60°C for 30s. Genome copy number was determined using a standard curve generated from serial dilutions of genomic DNA from purified EBs.

## ACKNOWLEDGEMENTS

We would like to thank Dr. Anthony Maurelli (University of Florida) and his laboratory for providing us with the strains used in this work as well as helpful feedback during the development of this project and drafting of the manuscript. We would also like to thank Dr. Pei Zhou (Duke University) for graciously providing us with the LpxC inhibitor used in this work. This work was supported by a MIRA ESI award (R35 GM138202) and a USU faculty start up award (HP73LIEC18) to GL. The funders had no role in study design, data collection and interpretation, or the decision to submit the work for publication. The opinions and assertions expressed herein are those of the author(s) and do not necessarily reflect the official policy or position of the Uniformed Services University or the Department of Defense.

## AUTHOR CONTRIBUTIONS

Conceptualization and Design, AD and GL; Data Curation and Formal analysis, AD and GL; Investigation, Methodology, Validation, Visualization, AD, GO, PS, and GL, Writing – original draft, GL; Writing – review & editing, AD, GO, PS, and GL; Funding acquisition, Project administration and Supervision, GL. All authors read and approved the final manuscript.

